# Mapping glioblastoma spreading: connexin43 and glial dynamic in mouse and human glioblastoma microenvironment

**DOI:** 10.64898/2025.12.16.694626

**Authors:** Assunta Virtuoso, Giampaolo Milior, Ciro De Luca, Julien Moulard, Luca Riccio, Alesya Evstratova, Anaïs Venard, Giovanni Cirillo, Johann Pallud, Gilles Huberfeld, Nathalie Rouach, Michele Papa

## Abstract

High-grade gliomas (HGGs), including astrocytoma and glioblastoma (GBM), constitute the most prevalent primary tumors of the central nervous system (CNS). GBM cells demonstrate a notable ability to infiltrate the brain parenchyma, precluding complete surgical resection. Here we investigated the spreading of GBM cells and the response of the CNS microenvironment focusing on glial cells, which are essential interactors to GBM.

We used acute and organotypic slices from the mouse brain and peritumoral cortex of patients with HGGs. We found that human peritumoral tissue from cortical resection was characterized by high levels of the astrocytic Connexin43 protein (Cx43) and discrete infiltration of microglia. In contrast, the tumor core exhibited high myeloid infiltration and an altered extracellular matrix (ECM) composition, which was poor in CD44. We tracked mouse and primary human-labeled-GBM cells in 2D cultures and in co-culture with organotypic slices generated from mouse brain and human peritumoral tissues. We found that the implanted GBM cells infiltrated the brain tissue, implying early glial modifications including an increase in Cx43 expression and distribution. Furthermore, the blockage of Cx43 hemichannels was accompanied by morphological changes and polarization of human GBM cells, typical for migration phenomena. The present study sheds light on the dynamics of GBM cells spreading in the living brain tissue, suggesting that the progression of the tumor correlates with changes within the host brain. Our findings identify the upregulation of Cx43 expression as a highly consistent modification in both mouse and human tissue that may be crucial for GBM infiltration.

## 1 Introduction

Diffusely infiltrating gliomas, including glioblastomas, astrocytomas, and oligodendrogliomas, are the most common type of primary brain tumors. Glioblastoma (GBM) cells have a remarkable capacity to diffusely invade the brain, making complete surgical resection impossible. Thus, the dispersion of GBM cells is a major obstacle for treating these tumors [1]. The patterns of GBM invasion are not random [2]. Rather, glioma cells tend to migrate with certain preferred paths such as along blood vessels or fiber tracts [3]. The majority of the GBM migrate along the corpus callosum and anterior commissure to reach the contralateral hemisphere [4,5]. To accomplish this, tumor cells squeeze through tight spaces of the brain parenchyma, modifying the microenvironment structure and functional networks.

All multicellular systems function through dynamic and coordinated action of cell circuits, changing their activity in response to environmental signals, with time stamps into cells and a plethora of time-dependent transcriptomic, signaling and cell-cell interaction [6]. Differentiated GBM cells, adherent or migratory, keep contact within the extracellular matrix (ECM) [7], integrate into the neuroglial circuits and modify their functional aspects [8,9]. Among the early interactions, the GBM deals with the brain innate immune system represented by astrocytes and microglia [10]. In vivo imaging showed that microglial cells sense the glioblastoma and become activated as soon as 30 min after the tumoral cells seeding [11]. Unlike the microglia, astrocytes are uniquely enriched in the peritumoral area [12] and form a sheet-like structure at the tumor edge, being physically connected with the tumor, in particular via the gap-junction protein Connexin43 (Cx43) [13,14]. As a consequence, electrical activity is altered and cortical plasticity is triggered [15,16]. Notably, GBM cells exhibit autonomous Ca^++^ oscillations that affect the network activity, altering the physiological brain rhythm and contribute to treatment resistance[17,18].

Astrocytes play a pivotal role in the GBM development, progression and therapeutic outcomes. Multiple signaling pathways have been described [19]. However, the role of glial Cx43 is still under debate. Cx43 is a multifunctional protein, with channel, hemichannel and non-channel functions. Cx43 is a ubiquitous protein, predominantly expressed by astrocytes, and it is involved in the formation of electrical synapses among the functions [20]. The channels are involved in cell-cell communication and are essential for brain homeostasis and calcium signaling [21]. The hemichannels (HCs) are instead involved in the autocrine and paracrine communication. GBM appears to release ATP, glutamate and extracellular vesicles (EVs) via the hemichannels for signaling to astrocytes, microglia, and other CNS cells [22–24]. Therefore, glial Cx HCs play a major role in purinergic signalling and ECM remodeling, tissue hyperexcitability and epileptogenesis [25–27]. The pharmacological inhibition of Cx HCs has been shown to alleviate neuroinflammation, blood-brain barrier dysfunction, and suppress seizure activity in animal models [28,29]. Furthermore, a monoclonal antibody that inhibits Cx26, Cx30 and Cx32 HCs has been demonstrated to be efficacious in both disrupting GBM progression and reducing hyperexcitability in mouse and human preclinical models [30]. The inhibition of Cx43 channel C-terminus has been implicated in sensitization to the chemotherapeutic agent temozolomide [31].

A role for Cx43 has been suggested in brain tumor invasiveness [32], as in a syngeneic intracranial mouse glioma model, Cx43-mediated intercellular communication between astrocytes is involved in the spreading of glioma cells into the brain parenchyma [33]. Yet whether Cx43 expression and distribution are regulated by GBM infiltration in the remodeled resident tissue is unknown. To investigate this possibility, we characterized the progression of mouse and eGFP-labeled human glioblastoma cells into mouse and human organotypic brain slices, respectively. The present study sheds light on the dynamics of glioma cell infiltration in living brain tissue and suggests that GBM progression induces structural modifications in the peritumoral microenvironment, with involvement of the Cx43. Blockade of Cx43 hemichannels induces morphological changes and polarization of human GBM cells. Altogether, these effects are similar in mouse and human tissues, suggesting conserved mechanisms across species and translational value of the present research.

## 2 MATERIALS AND METHODS

### 2.1 Murine Glioblastoma cell culture

The murine glioma GL261 cells were purchased from Creative Bioarray (cod. CSC-C9184W). GL261 cell line is a well-characterized and widely used model to investigate GBM cell biology, heterogeneity and immunotherapy [34]. GL261 cells were cultured in 90% Dulbeccós MEM + 10% h.i. FBS as suggested by the manufacturer, or in B12-enriched medium (Neurobasal-A, 2% B12 supplement, 1% N2 supplement, 1% glutamine, 0.5% glucose, 10 U/mL penicillin, and 100 μg/mL streptomycin) for the co-cultures with mouse organotypic slices [35].

### 2.2 Generation of eGFP Expressing Human Tumor Cells

Human primary tumor cells characterized as GBM cells were gently provided by GlioTex facility (Prof. A Idbaih and Dr. M Verreault). Human primary GBM cells were genetically modified to express eGFP using lentiviral vector. GBM cells were called as N2 and were transfected with the pEGFP-N1 expression vectors (Clontech), grown in selection medium containing G418 (500 lg/ml), and then FACS-sorted to select cells expressing high levels of eGFP. This method yielded populations of glioma cells in which greater than 95% of the cells expressed the fluorescent protein. N2 cells were grown in DMEM/F12 (Life Technologies,31331028), 100 UmL−1/0.1 mgmL−1 Pen-Strep, B27 supplement 50x (Life Technologies, 17504-044), Human FGF Basic (Peprotech, 100-18B), Human EGF (Peprotech, 100-15).

### 2.3 Animals

Young adult (3–5 week-old) C57Bl/6 mice (Envigo) (*n* = 30) were included in the present study. Animal care was in compliance with the Italian and European Guidelines for use and care of laboratory animals (EU Directive 2010/63) under the authorization no. 268/2012-B from the Minister of Health. Each animal was given free access to food and water, under a 12/12 h light/dark cycle.

### 2.4 Patients

The present study includes human brain tissues from 13 patients affected by glioma. Most cases were astrocytoma IDH1-mutant (R-132-H) (10/13; 77%), followed by oligodendroglioma IDH-mutant-1p/19q-codeleted (3/13; 23%). Most patients (11/13; 84.6%) had epileptic seizures at diagnosis. The samples were collected at Sainte-Anne Hospital and La Pitié-Salpêtrière Hospital (Paris, France). This study was approved by the Comité d’Evaluation et d’Ethique de l’INSERM - IRB00003888 (approval n°21-864). Subjects’ consent was obtained according to the Declaration of Helsinki.

### 2.5 Preparation of brain acute and organotypic slice cultures from young adult mouse

Animals were rendered unconscious using isoflurane gas, and then euthanized by cervical dislocation. Brains were dissected and coronal slices (300 µm) were cut using a vibratome (Campden Instruments, 5100mz), while maintaining the sterility, as previously reported [36]. Mouse slices to culture were transferred in 70 µm cell strainer with fresh, RT carbogen-oxygenated ACSF (artificial cerebrospinal fluid) for recovery (45 min). Slices were cultivated in B12-enriched medium, as per the GL261 cell culture. Brain tissue cultures were maintained in a humidified atmosphere with 5% CO2 at 35°C. The medium was replaced three times a week. The young adult mouse organotypic slices were cultured up to 7 DIV.

### 2.6 Preparation of human acute and organotypic slice cultures

Surgical specimens were obtained from patients with glioma. During surgery, a small block of cortical tissue was removed as a security margin of glioma supratotal surgery approach, and was immediately placed in ice-cold sucrose ACSF and transferred to the laboratory. The tissue was processed as previously described [37]. Briefly, the tissue block was immerged in carbogen-oxygenated ACSF, cleaned of blood clots, vessels and meninges, and cut in 350 µm thick slices using the vibratome (Leica VT1200) at low speed (0.1 mm/sec). Each acute slice to be processed was immediately transferred to the interface chamber for recordings, while others were fixed in Zinc Formalin Fixative (ZFF) (Sigma-Aldrich, Z2902).

Human slices to culture were transferred in cell strainers with fresh, RT carbogen-oxygenated ACSF for recovery (45’min). After that, they were cultured for 1 h in NCS medium, in incubator with 5% CO2 at 37°C. The NCS is composed by 50% Neurobasal-A, 50% DMEM, Hepes 20mM, 2% B27 supplement, 1% N2 supplement, 1% Glutamax, 100 UmL−1/0.1 mgmL−1 Pen Strep, 1% NEAA. The NCS medium was replaced by fresh human cerebrospinal fluid (hCSF), which was changed three times per week. The human organotypic slices were cultured up to 7 DIV.

### 2.7 Viability test for GBM cells and organotypic slices cultures

Cell viability was assessed using 3-[4,5-dimethylthiazol-2-yl]-2,5 diphenyltetrazolium bromide salt (MTT) (Sigma-Aldrich), as previously described [38]. Briefly, GL261 cells were seeded at a density of 1 × 10^4^ cells/well in 100 μL of B12-culture medium, in 96-well plates. At confluence, the medium was replaced with fresh medium with MTT salt added to each well for 4 h at 37°C. 100 µL of Dimethyl sulfoxide (DMSO) were used to dissolve formazan salts for 15 min at 37°C, and absorbance was measured at 570 nm in a plate reader (Parkinelmer, Victor Nivo). Three replicate wells were used for each group. The DMSO was used as a blank; GL261 treated with 10mM H_2_O_2_ were used as a (negative) control.

Organotypic brain slice cultures were incubated in sterile dPBS with MTT substrate for 20 minutes at 37°C. To evaluate the impact of PBS incubation on slice viability, an additional fresh brain slice in either sterile dPBS or dPBS containing 10 mM H₂O₂ (control) was included in the analysis. Afterwards, brain slices were transferred into fresh tubes and DMSO was added in a volume proportional to the slice weight to dissolve the violet formazan crystals formed during the assay. The samples were then centrifuged and 100 µL of the supernatant was transferred to a 96-well plate in duplicate. Absorbance was measured at 570 nm.

### 2.8 GBM cells injection in human and mouse organotypic slice cultures

The number of living and dead cells in culture was determined by cell counting after trypan blue staining as previously described [39]. Briefly, cells were dissociated with trypsin digestion. Cells suspension was diluted 1:1 in trypan blue solution (Sigma Aldrich), incubated for 2 min and counted using a Burker chamber. Cell viability was determined based on the total cell count, the dilution factor and the trypan blue dye exclusion assay. For human and mouse cell transplants, N2 and GL261 cells were centrifuged and resuspended in PBS (10000 cells in 1 µL of PBS). They were immediately injected into the human cortical slices or cortex from whole brain mouse slices respectively, using a sterile patch-clamp tip connected to a syringe. GBM cells engrafted the slices and their integration could be observed in the following days as previously described [40].

### 2.9 Drug Treatments

The synthetic peptides TAT-GAP 19 (>85% pure) was obtained from GenScript (Piscataway, NJ, USA). YGRKKRRQRRR was the TAT sequence, which is responsible for the cell penetration of the peptides [41]. Gap19 is a selective inhibitor of the Cx43 hemichannels (HC). It perturbs the carboxy terminal interaction with a cytoplasmic loop (CL)-located binding site, leading to the Cx43 HC closure [42].

GAP-19 was dissolved in sterile ddH20 as per manufacturer’s instructions, at a final concentration of 200 µM, and added to the culture medium. Control peptide was administered in an identical manner. The half-life of Tat-Gap19 peptide was calculated with the peptide lifetime predictor (PlifePred, http://crdd.osdd.net/raghava/plifepred/) as described in [43].

GAP-19 peptide was provided to GBM cells or to human organotypic slices with or without GBM cells injection. The potential effects of GAP-19 on treated, untreated GBM cells, and human organotypic slices were investigated using dynamic microscopy. GBM cells and slices were fixed at 4h from the last treatment and prepared for immunostainings.

### 2.10 Scratch and invasion assay

To evaluate the effects of GAP19 on the migratory properties of GBM cells, GL261 cells were cultured on 12-well plates until they reached 90% confluence. After a gentle scratch, GBM cells were washed twice with culture medium to remove cell debris and incubated with Gap-19 200 µM until endpoints (4 h and 24 h). Images were acquired with a IXplore IX73 Standard microscope, Evident. The distance between the two edges of the scratch was labeled and measured. Migrating cells were defined as cells found into the scratch. Cell migration ability was evaluated by counting the cells into the scratch in at least 4 random fields/well.

### 2.11 Ex Vivo electrophysiology: Multi Electrode Array (MEA) Recordings

Human acute and organotypic slices were transferred in an interface chamber containing a standard artificial cerebrospinal fluid (ACSF; 119mM NaCl, 2.5mM KCl, 2.5mM CaCl_2_, 1.3mM MgSO_4_, 1mM NaH_2_PO_4_, 26.2mM NaHCO_3_, and 11mM glucose, saturated with 95% O_2_ and 5% CO2, at a temperature of 37°C) for at least 1h. For multi-electrode array (MEA) recordings, slices were transferred on planar MEA petri dishes (200-30 ITO electrodes, organized in a 12×12 matrix, with internal reference, 30μm diameter and 200μm inter-electrode distance; Multichannel Systems, Germany). They were kept in place by using a small platinum anchor. The slices on MEAs were continuously perfused at a rate of 2 mL/min with a magnesium-free ACSF. Pictures of slices on MEAs were used to identify the location of the electrodes through the different cortex regions and to select the electrodes of interest. Data were sampled at 10kHz and network spontaneous activity was recorded at room temperature by a MEA2100-60 system (bandwidth 1-3000Hz, gain 2x, Multichannel Systems, Germany) through the MC Rack 4.5.1 software (Multichannel Systems, Germany) [44]. Slices were then perfused with an “epileptogenic” ACSF, in which Mg^2+^ is absent and K^+^ is raised to 6 mM, to boost tissue excitability. Once the perfusion with 0 Mg^2+^ 6 mM K^+^ ACSF had started, the activity of cortical slices was recorded.

### 2.12 Dynamic Microscopy of GBM cells and Brain Slice Cultures

Dynamic photography was performed by taking a picture of the cells and slices from a single focal plan every 3 hours over a time frame of 7 days under a fluorescent, inverted microscope (Thermo Fisher Scientific, EVOS Microscope). N2 morphometry was quantified using ImageJ software (Rasband, W.S., ImageJ, U. S. National Institutes of Health, Bethesda, Maryland, USA, http://imagej.nih.gov/ij/, 1997–2011).

Mouse and human acute/organotypic slices were fixed in 4% paraformaldehyde or zinc formalin, respectively, at 3 and 7 days post-injection (dpi). The immunostaining protocol was adjusted from previous publications [25,45–47]. Briefly, the slices were incubated for blocking in 1% Tryton, 10% Serum in Pbs/NaN_3_ and then with primary antibodies for 4 overnights (see **Table 1**). After washing in PBS, 3 over night incubation with the appropriate secondary antibodies (Alexa Fluor) and Dapi (4’,6-diamidino-2-phenylindole) nuclear staining was performed. Tissues were imaged using a laser scanning confocal system microscope (Zeiss LSM 900). Two and three-dimensional reconstructions of brain tumor slices were generated from fluorescent micrographs.

**Table 1.** Primary Antibodies used.

| Target | Brand | Dilution |
| --- | --- | --- |
| anti-GFAP | AB5804, Millipore | IF: 1:400 |
| anti-Iba1 | 019-19751, WAKO | IF: 1:500 |
| anti-Connexin43 | A1978, C6219, Sigma-Aldrich | IF: 1:200; WB:1:1000 |
| anti-Ki67 | ab16667, Abcam | IF: 1:50 |
| anti-CD44 | Ab189524, Abcam | IF:1:200 |
| Anti-β-actin | A5441, Sigma-Aldrich | WB: 1:10000 |

GBM cell cultures were stained with a similar protocol, reducing incubation times (1 overnight for the primary antibodies, 2h for the secondary antibodies), and imaged using an inverted fluorescence microscope (Thermo Fisher Scientific, EVOS Microscope or IXplore IX73 Standard microscope, Evident).

### 2.13 Western Blotting

Brain slices protein extraction was performed by adding 100µL of RIPA buffer (140 mM NaCl, 10 mM Tris-HCl pH 8.0, 1 mM EDTA, 1% Triton X-100, 0.1% SDS, 0.1% sodium deoxycholate, 1 mM PMSF) supplemented with fresh protease and phosphatase inhibitors (1% each). Samples were vortexed on ice for 5 minutes and homogenized using a loose pestle. Homogenates were centrifuged at 14,000 rpm for 10 minutes at 4 °C to separate the supernatant and pellet. The pellet was resuspended in 50µL of RIPA buffer. Protein concentrations of both supernatant and resuspended pellet were determined using a Bradford assay in 96-well plates and measured with a VICTOR Nivo plate reader (Revvity Life Sciences).

Samples were prepared for SDS-PAGE by mixing with 4× Laemmli Sample Buffer (60 mM Tris-HCl pH 6.8, 10% glycerol, 2% SDS, 100 mM DTT, 0.1% bromophenol blue) to a final volume of 15µL containing 30µg of total protein. Proteins were resolved by SDS-PAGE using a discontinuous polyacrylamide gel system (stacking and resolving). The stacking gel (5% acrylamide/bis-acrylamide) contained 137 mM Tris-HCl pH 6.8, 0.1% SDS, 0.1% APS, and 0.1% TEMED. The resolving gel (12% acrylamide/bis-acrylamide) contained 300 mM Tris-HCl pH 8.8, 0.1% SDS, 0.1% APS, and 0.05% TEMED.

After electrophoresis, proteins were transferred onto a nitrocellulose membrane using electroblotting. The transfer was carried out in Transfer Buffer (0.24 M Tris, 0.2 M glycine, 20% methanol, pH 8.1–8.4). Successful transfer and uniformity were verified by staining the membrane with 0.1% Ponceau Red in 5% acetic acid, followed by washes in TBST (10 mM Tris-HCl pH 7.5, 150 mM NaCl, 0.1% Tween-20).

Membranes were blocked in 5% BSA in TBST and incubated with primary antibodies targeting proteins of interest (see **Table 1**). After overnight incubation, membranes were washed three times in TBST and then incubated with species-specific HRP-conjugated secondary antibodies. Following secondary antibody incubation, the same washing steps were repeated.

Detection was performed using enhanced chemiluminescence (ECL) solution containing luminol and hydrogen peroxide, substrates for horseradish peroxidase (HRP). Membranes were incubated with equal volumes of ECL components for 1 minute, and chemiluminescent signals were captured using a LI-COR Digit imaging system (6-minute exposure). Densitometric analysis of protein bands was conducted using ImageJ software.

### 2.14 Measurements and Statistics

Raw data from MEA recordings were analyzed using MC Rack (Multi-Channel System, Reutlingen, Germany). Detection of bursts was performed using the “Spike Sorter” algorithm, which sets a threshold of the noise (5-fold), and low-pass filter. This allowed to automatically detect each event [44]. Measurements of immunofluorescence (IF), and western blotting (WB) markers were performed using ImageJ software (Rasband, W.S., ImageJ, U. S. National Institutes of Health, Bethesda, Maryland, USA, http://imagej.nih.gov/ij/, 1997–2011) in a blinded manner.

Data were exported and analyzed using the Graph Pad Prism 8.4 program (SPSS-Erkrath). Data were checked for normal distribution and homogeneity of variance by the Kolmogorov–Smirnov’s and Levene’s mean tests, respectively. Normal distributions with equal variances were analyzed by Student’s t tests for comparing two groups, and one-way ANOVA for multiple comparisons followed by all pairwise Holm–Sidak post hoc test. Kruskal–Wallis test was used for non-parametric analysis followed by Tukey’s test for pairwise multiple comparisons. Level of significance was set at p = 0.05 (*p ≤ 0.05; **p ≤ 0.01; ***p ≤ 0.001). Statistical techniques were employed to derive correlation coefficients and mutual information metrics for each feature. Results are expressed as group mean ± SEM and represented in the charts.

## 3 RESULTS

### 3.1 Characterization of Glioblastoma cells

We characterized GL261 cells and tested their viability in B12-enriched medium to obtain optimal murine brain slice-tumor co-cultures. GL261 cells have been characterized in detail [34]. They appear with the shape of fibroblastoids, but have a genetic profile similar to the primary human GBM, making them a very aggressive tumor. By reaching confluence, they form adherent neurospheres. In the B12-enriched medium they retain the morphology, arrange in neurospheres, do not express GFAP and are viable at higher rates compared to the standard medium (One-way Anova, Tukey’s multiple comparison test, F (2,17), df=2, p<0.0001) (**Fig.1A-C**). These results indicate that GL261 do not show gross changes in the enriched microenvironment, compared to the basic medium suggested by the manufacturer.

**Fig. 1.**
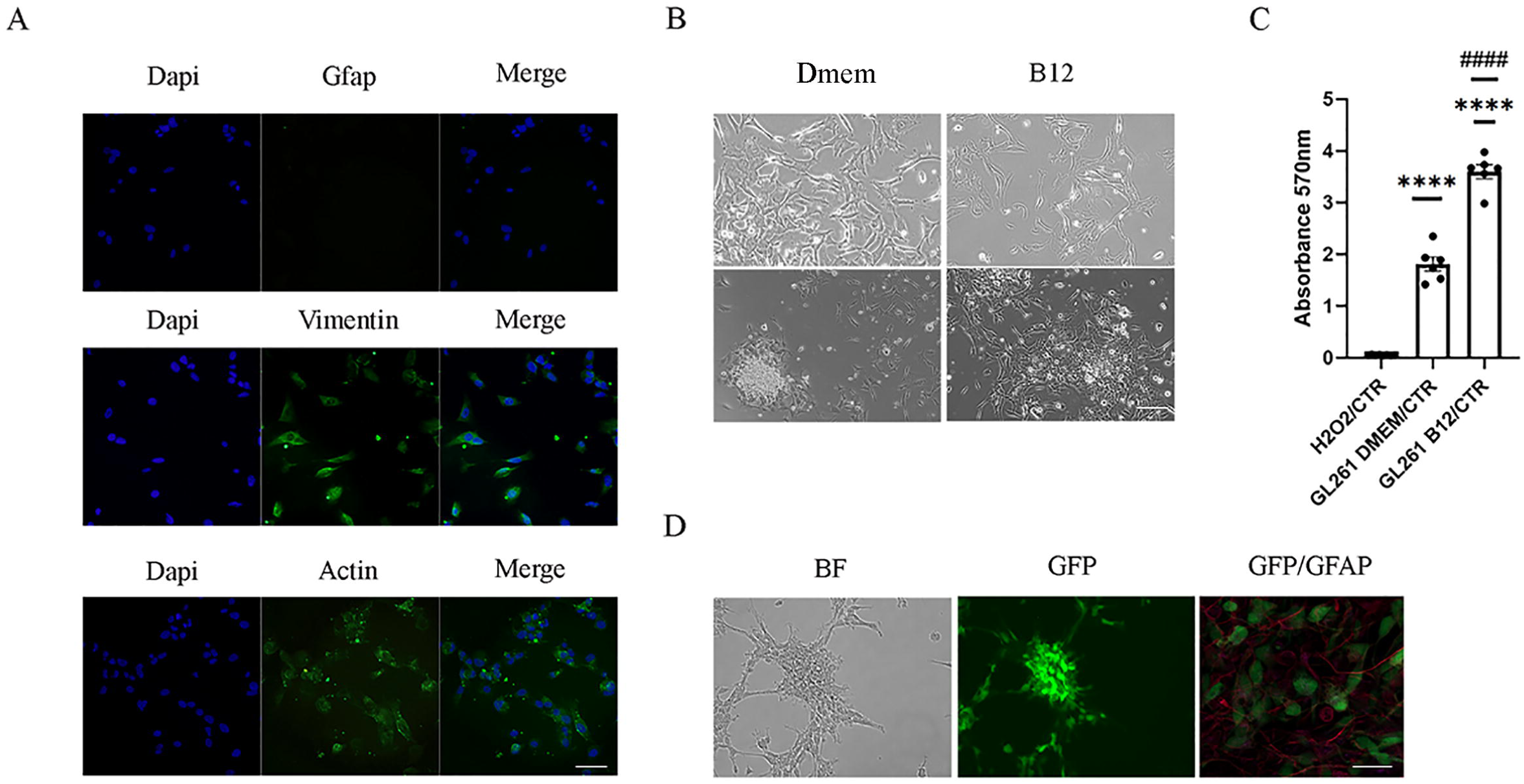
Characterization of GBM cells. **(A)** Representative immunofluorescence imaging for structural markers in murine GL261 cells, GFAP, Actin and Vimentin (green) in B12-enriched medium. Nuclei were stained using DAPI. Scale bar 50 µm. **(B)** Morphological features of GL261 cells grown in DMEM/10%FBS (Dmem) and B12 enriched medium (B12). Scale bar 50 µm. **(C)** Viability test using MTT substrate for GL261 cells grown in DMEM/10%FBS and B12 enriched medium (One-way Anova, Tukey’s multiple comparison test, F (2,17), df=2, p<0.0001) **(D)** Morphological features of human primary GFP^+^-N2 cells. BF=bright field; GFP, green=green fluorescent protein; GFAP, red= Glial fibrillary acidic protein. Scale bar 100 µm.

Human primary N2 cells were cultured in DMEM/F12 supplemented with B27 1x, EGF 20ng/mL and b-FGF 20ng/mL. N2 cells have a similar aspect compared to the murine GL261 cells. N2 cells form neurospheres. However, they express GFAP (**Fig.1D**).

### 3.2 The neuroglial network changes among tumor and peritumoral tissues

A macroscopic evaluation was conducted on the tissue collected from patients affected by high grade glioma. We classified the tissue pieces in “tumoral” or “peritumoral” based on the technical information delivered by the neurosurgery unit. We confirmed the macroscopy of healthy tissues for the peritumoral samples. To study the modifications of the human tumor microenvironment, we performed immunostainings in combination with antibodies that selectively label astrocytic-like cells (anti-Cx43), myeloid cells (anti-Iba1), and the extracellular matrix receptor (anti-CD44), which has been proposed as an invasion marker for GBM [48].

Cx43 has a dynamic expression. It can be found with a dispersed cytoplasmic, perinuclear or membrane localization [49]. However, the autofluorescence of the lipofuscin granules may interfere with the perinuclear targeting of the marker. For accurate analysis, we considered the mean density value related to the area of interest, with spatial exclusion parameters. Peritumoral pieces showed a high density for Cx43 compared to the tumoral pieces (p= 0.0366, two-tailed unpaired t-test with Welch’s correction, t=2,282, df=15,90) (**Fig. 2A**). Cx43 expression correlates with CD44 levels during the GBM invasion [23]. Accordingly, we found a spatial, positive molecular distribution among the markers. Peritumoral acute slices showed a high CD44 density compared to the tumoral samples (p= 0.02, two-tailed unpaired t-test, t=2,643, df=13) **(Fig. 2 B)**.

**Fig. 2.**
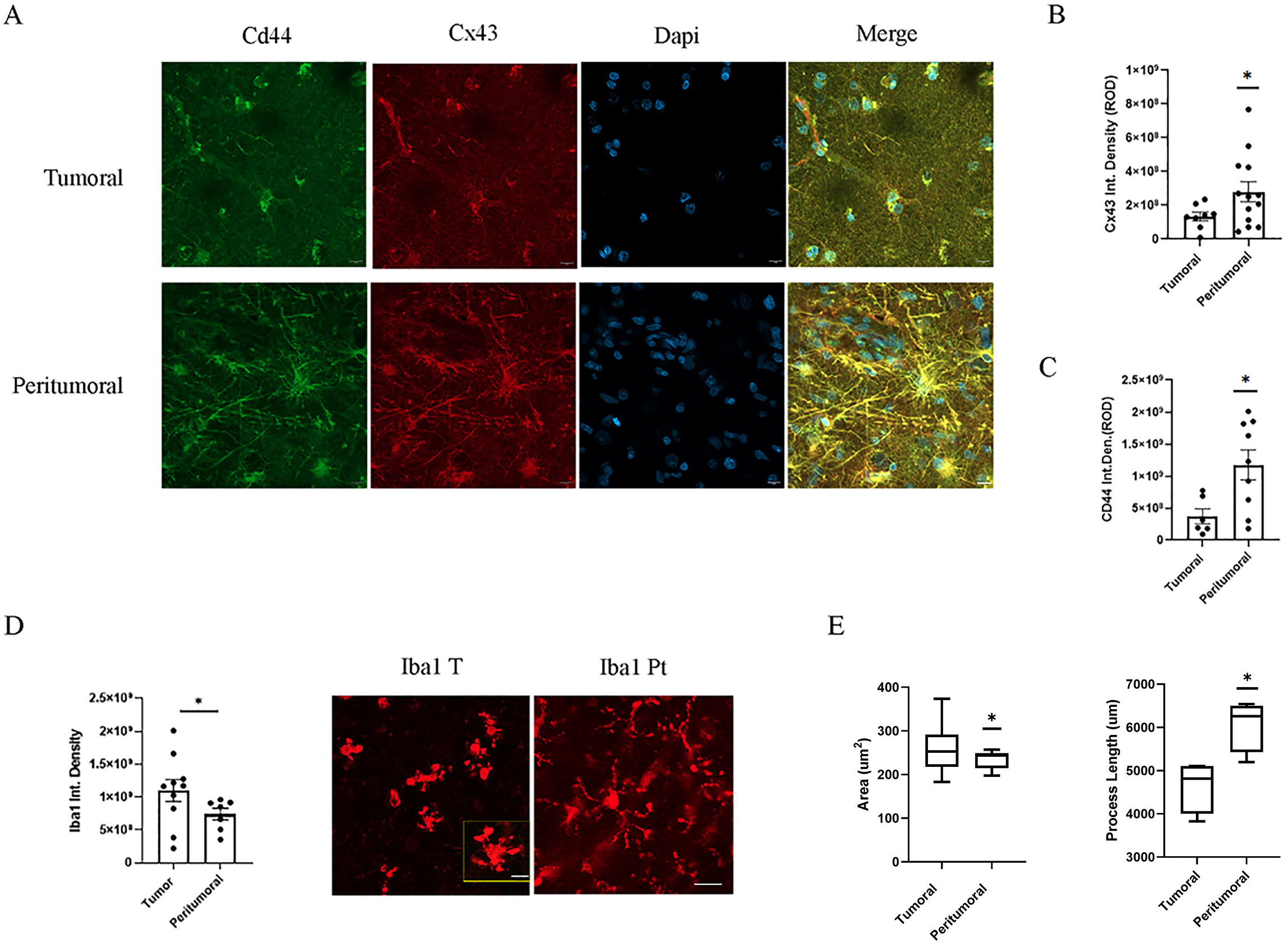
Characterization of acute human tissue microenvironment from patients with glioma. **(A)** Representative immunofluorescence for Cx43 and Cd44 images acquired by using the confocal microscopy of acute tissue slices among tumoral and peritumoral tissue. Nuclei were stained by DAPI (blue). Scale bar 20µm. (**B**) Relative quantification of Cx43 intensity from confocal imaging of acute tissue slices among tumoral and peritumoral areas (p=0.0366, two tailed unpaired t-test with Welch’s correction, t=2.282, df=15.90). (**C**) Relative quantification of CD44 intensity from confocal imaging of acute tissue slices among tumoral and peritumoral tissue (p= .0002, two tailed unpaired -est, t=5.818, df=10). (**D**) Representative immunofluorescence imaging for Iba1 positive cells from confocal imaging of acute tissue slices among tumoral and peritumoral areas, and relative quantification of signal intensity (p=0.021, Kolmogorov-Smirnov test). Scale bar 50µm; 100 µm for the inset magnification. (**E**) Morphometric measurement of Iba1+ cells from confocal imaging of acute tissue slices among tumoral and peritumoral areas (mean cell area, p=0.0250, two tailed unpaired t-test with Welch’s correction, F test to compare variances t=1.120, df=12), (processes length, p=0.0058, two tailed unpaired t-test, t=3,329, df=6).

Immunostaining analysis for the microglial Iba1 marker revealed that Iba1 density was higher in tumor pieces compared to the peritumoral tissues (p= 0.021, Kolmogorov-Smirnov test) **(Fig. 2 C)**. Myeloid cells from the tumor core showed an increase of the area and a reduction of the mean processes length, typical of the macrophagic shape (regarding mean cell area, p= 0.0250, two tailed unpaired t-test, t=1.120, df=15), (regarding processes length, p= 0.0058, two tailed unpaired t-test, t=4.608, df=5) **(Fig. 2 D)** [5].

This evidence proves that the glial microenvironment in human high-grade gliomas and their peritumoral tissues potentially has a distinguished morphological and molecular profile, which could be crucial for disease predictions, modeling and interventions.

### 3.3 Functional neuroglial networks in organotypic cultures

It is known that organotypic culturing modifies the neuroglial networks and the adult tissue is fragile [50]. We verified the vitality of our model by recording the electric activity of organotypic slices, with or without the GBM cells transplantation. We compared the organotypic recordings to acute conditions. The activity of human acute cortical slices gradually increases according to Dossi E et al, [37].With epileptogenic “ACSF” (6mM K^+^, 0 mM Mg^2+^), acute cortical slices showed a significant increase in the discharging frequency (p=0.0225, F test for variance, df=3.3) with a tendential reduction in the amplitude of the peaks **(Fig. 3A-C)**.

**Fig. 3.**
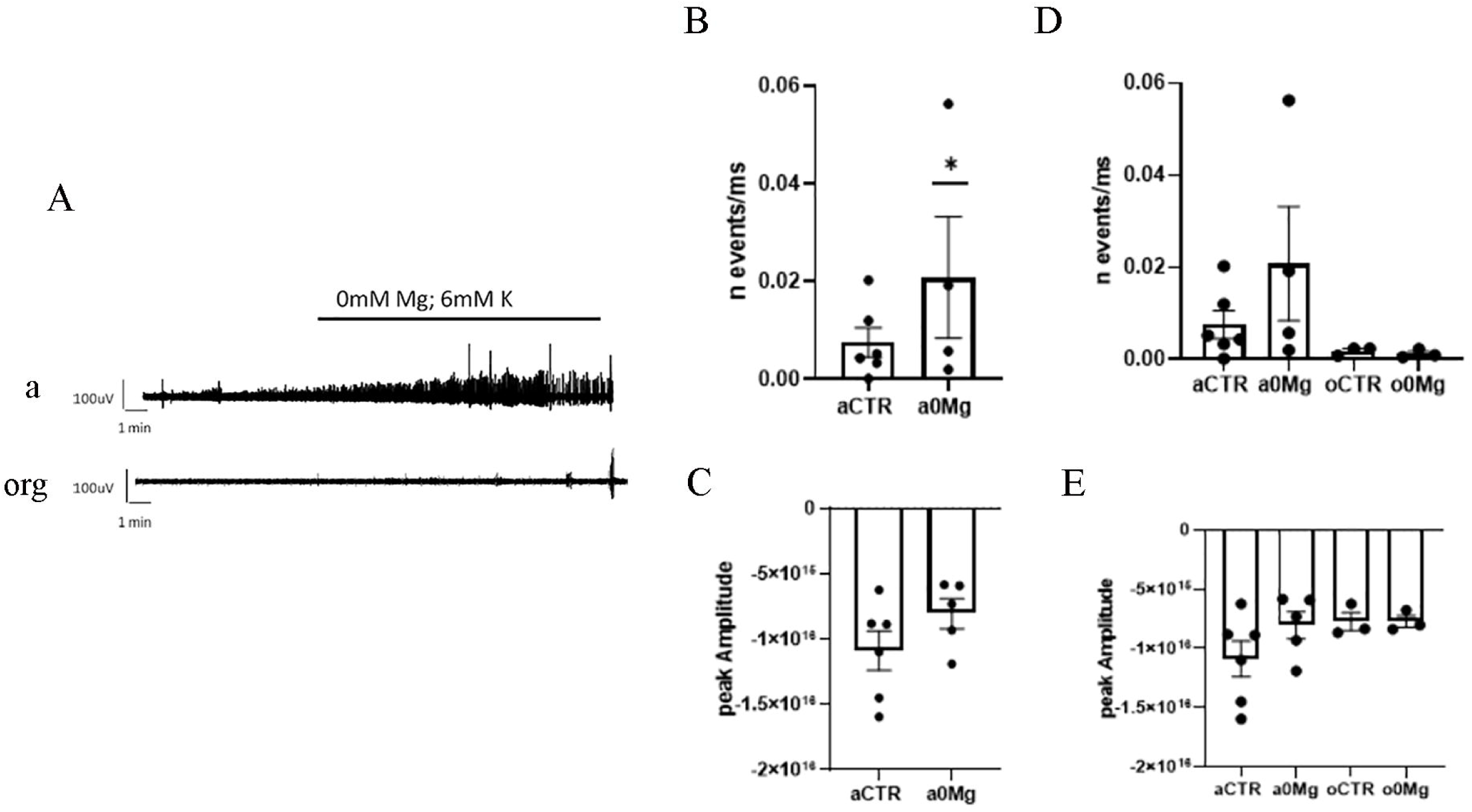
Electrophysiological characterization of acute human tissue from patients with glioma, and organotypic conditions. (**A-B**) Representative panel and analysis of frequency of electrical spikes in human acute slices. a, acute; org, organotypic. (**A-C**) Representative panel and analysis of amplitude of electrical spikes in human acute slices. (**A-D**) Comparative analysis of frequency of electrical spikes in acute and organotypic conditions. (**A-E**) Comparative analysis of amplitude of electrical spikes in acute and organotypic conditions.

Human organotypic slices with or without GBM cells transplantation were tested using MEA after 7 DIV. They appeared alive with a multiscale-remodeling of the circuits. Frequency of the events was much lower compared to acute conditions, while the amplitude of the peaks remained unchanged **(Fig. 3A, D-E)**.

These results indicate that our organotypic model maintained the vitality despite modifications of the neuroglial circuit.

### 3.4 Transplanted Glioblastoma Cells Infiltrate the Organotypic Brain Slices

To characterize the GBM progression and microenvironment, we injected mouse GL261 cells and human e-GFP+ GBM cells in the organotypic cortical tissue at div 1 in murine and human slices, respectively.

The adult mouse organotypic slice cultures almost retained their vitality for one week when injected with PBS compared to the fresh acute tissue (Acute 0.2906 ±0.016; Org PBS 0.2233±0.019). The young adult mouse cultures represented a fertile platform for the GL261 progression, as the vitality of the system triplicated (0.5516±0.038, p≤0.0001, One-way Anova followed by Tukey’s test for multiple comparisons) (**Fig 4A**). Accordingly, the proliferating cells expression marker ki67 was restricted to densely packed cells clusters with large nuclei (**Fig. 4B**).

**Fig. 4.**
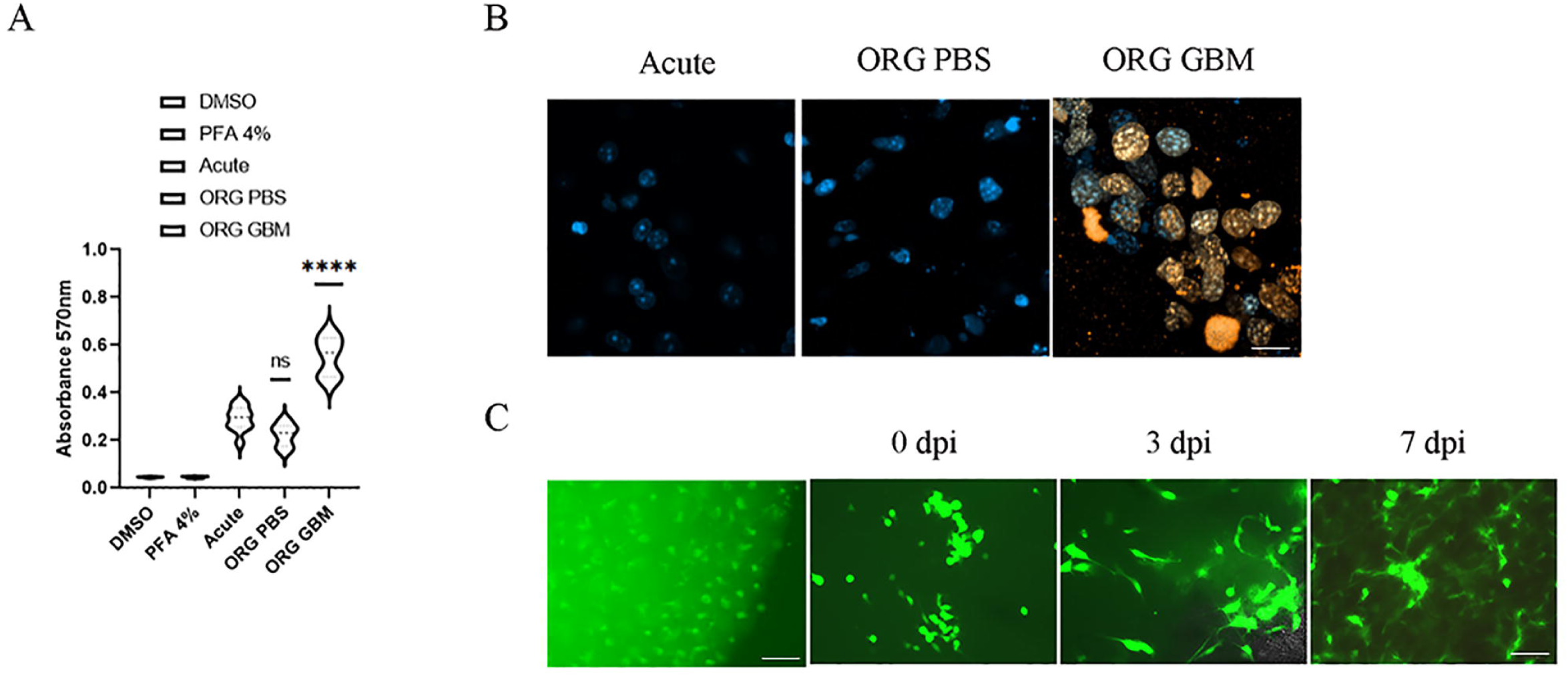
Glioblastoma infiltrates the microenvironment in the organotypic models. (**A**) MTT viability test for mouse organotypic slice cultures in acute and organotypic condition, with PBS or GBM injection. DMSO=CTR, PFA 4%=Negative CTR (p≤0.0001, One-way Anova followed by Tukey’s test for multiple comparisons). (**B**) Representative panel for immunofluorescence of the proliferative marker ki67 (orange) in acute and organotypic cultures of murine slices, with PBS or GL261 cells injection. Nuclei are stained by DAPI (blue). Scale bar 20µm. (**C**) Representative dynamic microscopy of transplanted eGFP-labeled GBM cells at 1, 3, and 7 dpi in human cortical tissue. The dark field image at 3dpi (left) was overlapped to the corresponding bright field in order to show the tissue in background. Scale bar 100 µm.

We examined the distribution of transplanted eGFP-labeled GBM cells at 1, 3 and 7 dpi. eGFP-labeled GBM cells were localized either in multicellular clusters or as individual cells. GBM cells had a predominantly unipolar morphology, with a rounded cell body in the first dpi. e-GFP GBM cells appear to increase the branching in the following days and completely cover the tissue, both in the gray and white matter (**Fig 4 C**), suggesting a spreading infiltration [1].

### 3.5 Glioblastoma Spreading modifies the microenvironment

To study the modifications of the glial microenvironment following GBM infiltration, we investigated the expression of the markers Cx43, GFAP, and Iba1 in both mouse and human organotypic slices injected with PBS or tumor cells.

Immunostainings for GFAP showed an increase in the fluorescent intensity in response to the murine GBM development comparing to the slices injected with PBS, suggesting a reaction of the resident astrocytes by 3 dpi (unpaired t’ student test, p=0.0087, t=3.340; df=9; One-way Anova followed by Tukey’s test for multiple comparisons, Acute vs Org PBS, p 0.0001; Acute vs Org GBM, p 0.015) (**Fig. 5A-B**). We found an increase in Cx43 protein expression in organotypic slices injected with GBM compared to the PBS delivery at 3 dpi (t student’ test, p value=0.055, t=3.961, df=7)(**Fig. 5B-C**). The increase in Cx43 expression may reflect a shift between its cytoplasmic and membrane form. Western blot analysis on soluble and insoluble tissue fraction showed that the co-culture GBM-brain slice undergo an increase of the ratio Cx43 Cytoplasm/Membrane compared to the acute system or to the GL261 2D cultures (One way Anova followed by Kruskal-Wallis test, Org GBM vs Gl261, p= 0.053; Org GBM vs Acute, p=0.2277) **(Fig 5D-E)**.

**Fig. 5.**
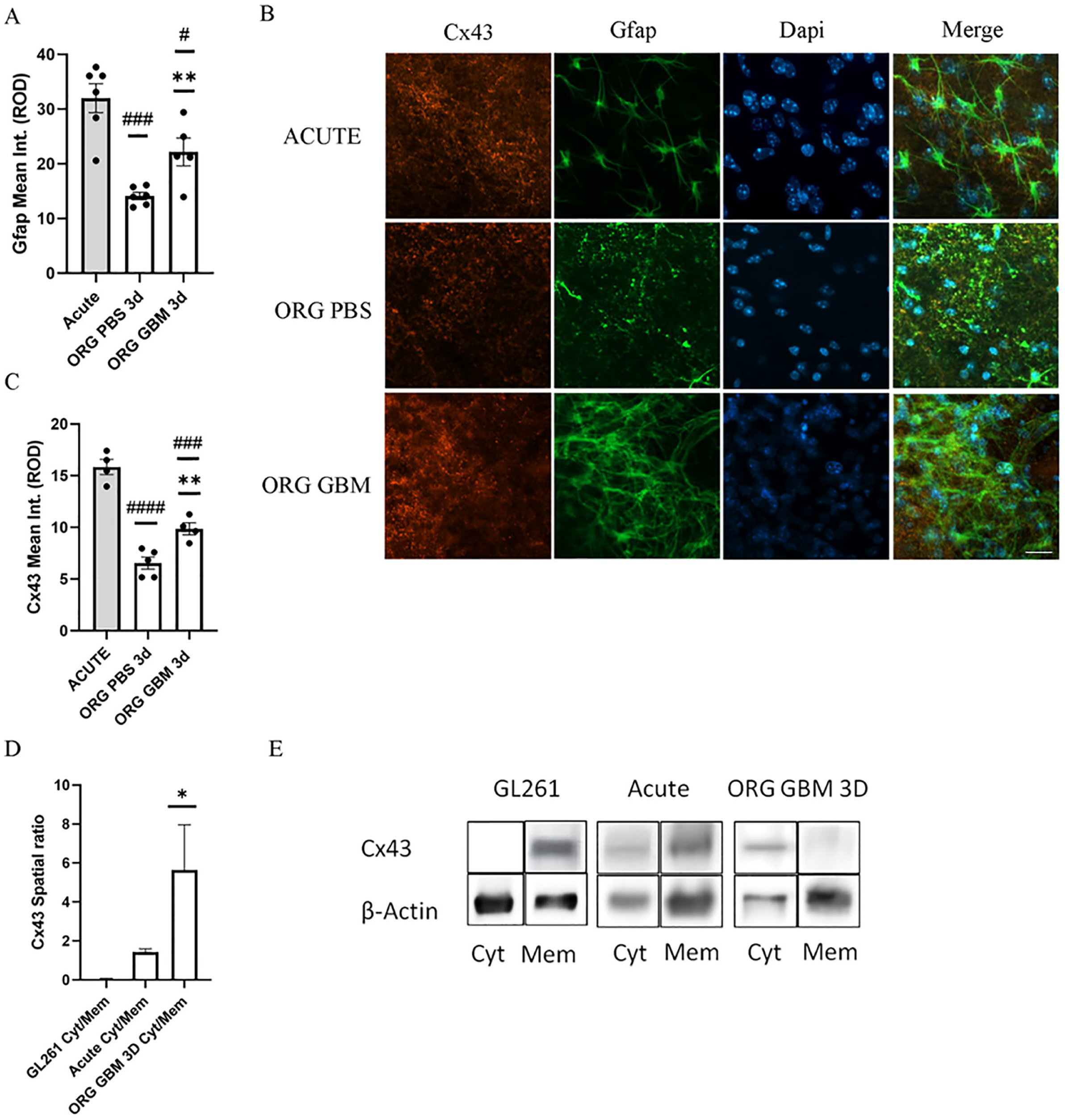
Murine Glioblastoma Infiltration modifies the microenvironment. (**A**) Analysis of GFAP fluorescence intensity in acute and organotypic slices, following PBS or GBM injection. (p>0.05, One-way Anova followed by Tukey’s test for multiple comparisons). (**B**) Representative immunofluorescence for Gfap and Cx43 images acquired by using the confocal microscopy of acute and organotypic tissue slices, following PBS or GBM injection. Nuclei were stained by DAPI (blue). Scale bar 20µm. (**C**) Analysis of Cx43 fluorescent signal intensity in acute and organotypic slices, following PBS or GBM injection. (p>0.05, One-way Anova followed by Tukey’s test for multiple comparisons). (**D**) Analysis of Cx43 protein localization ratio between the soluble (cytoplasmic) and insoluble (membrane) part from western blotting samples of mouse acute slices, 2D cultures of GL261 cells, and organotypic co-cultures (p>0.05, One-way Anova followed by Tukey’s test for multiple comparisons). (**E**) Representative western blotting bands from acute slices, 2D cultures of GL261 cells, and organotypic co-cultures from soluble (cyt, cytoplasmic) and insoluble (mem, membrane) part of the tissues.

Immunostainings in the human GBM-brain co-cultures system revealed an increase of expression (p=0.2929, t test, df=8)(data not shown) and cell number (p= 0.0437, t test, df=10) positive to Cx43 at 3 dpi **(Fig. 6 A-B)**. However, eGFP-labeled GBM cells expressed Cx43 at low levels, either in 2D culture or in co-culture with human tissue (**Fig. 6 A-C**). These results indicate that resident Cx43 is involved in the CNS response to the GBM progression. Accordingly, it has been shown using spatial transcriptomics that the invasive pattern of mesenchymal-like humanGBM may mimic an astrocytic-like microenvironment [51].

**Fig. 6.**
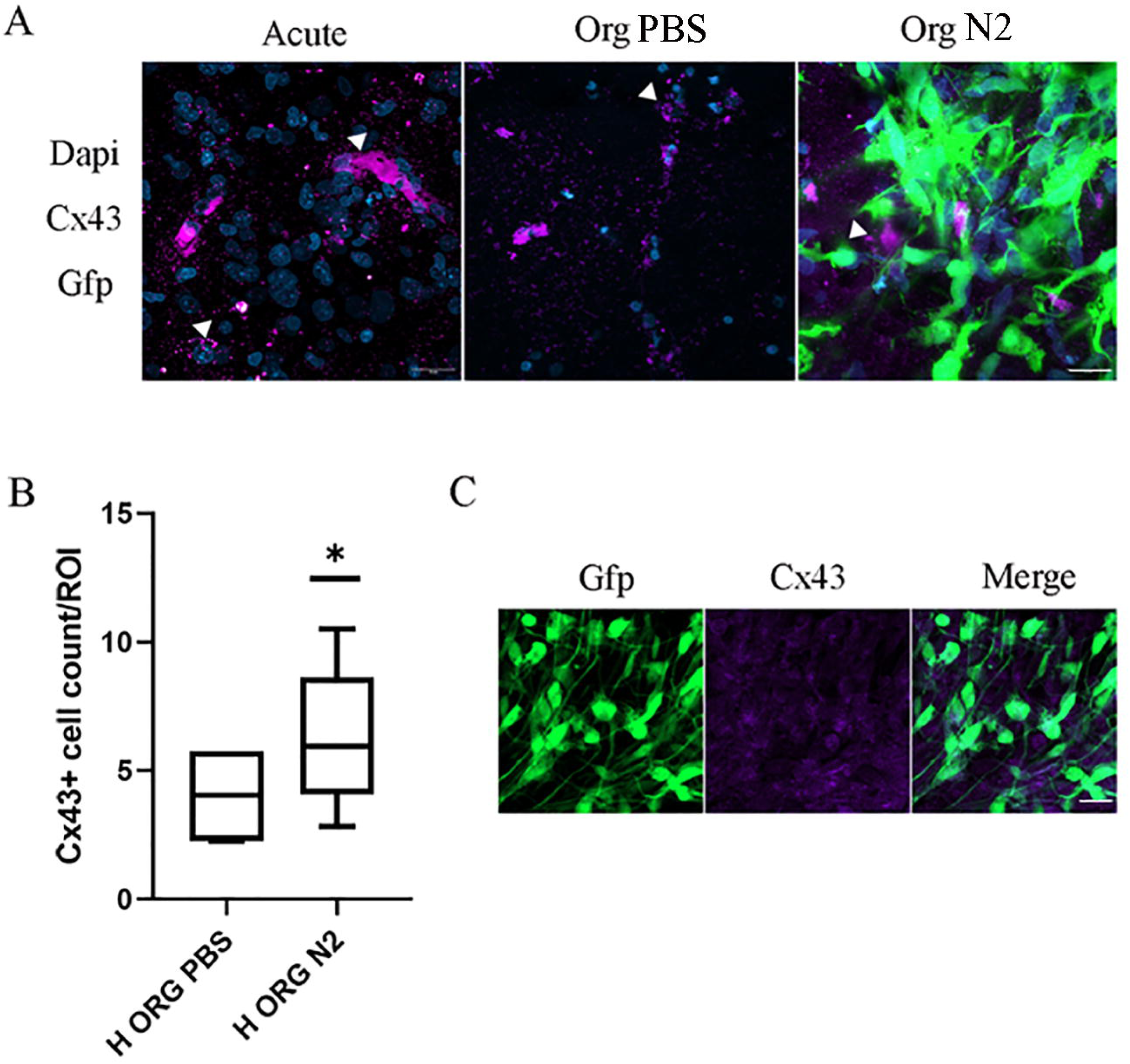
Glioblastoma modifies the human astrocytic microenvironment. **(A)** Representative immunofluorescence imaging for Cx43 positive cells (white arrows) from confocal imaging of organotypic tissue slices in CTR group or injected with PBS and/or N2 GFP+ cells. GFP, green; Cx43, purple; DAPI staining for nuclei, blue. Scale bar 20 µm. (**B**) Relative quantization of Cx43 cell count per region of interest (ROI) in organotypic tissue slices injected with PBS or N2 GFP+ cells (t-student test, expression, p=0.2929, t test, df=8; cell number p= 0.0437, t test, df=10). (**C**) Representative immunofluorescence imaging for human primary N2 cells. GFP, green, Cx43, purple. Scale bar 20 µm.

The myeloid component was affected by the organotypic culturing (**Fig 7 A-C**). We found a basal decrease of Iba1 expression in the organotypic models compared to the general acute conditions (p= 0.0417, one-way anova, Tukey’s multiple comparisons test, df=12) (**Fig. 7A-B**). Iba1-expressing cells indeed appeared to modify their area and reduce the length of the processes in the organotypic co-culture with GBM cells at 3 dpi (regarding area, Ordinary One way Anova followed by Tukey’s multiple comparisons test, acute vs org pbs, p=0.0334; org pbs vs org N2, p=0.0323; F=5.557, df=2), (regarding processes length, Ordinary One way Anova followed by Tukey’s multiple comparisons test, acute vs org pbs, p≤0.0001; acute vs org gbm, p≤0.0001; F=43, df=2) (**Fig. 7C**).

**Fig. 7.**
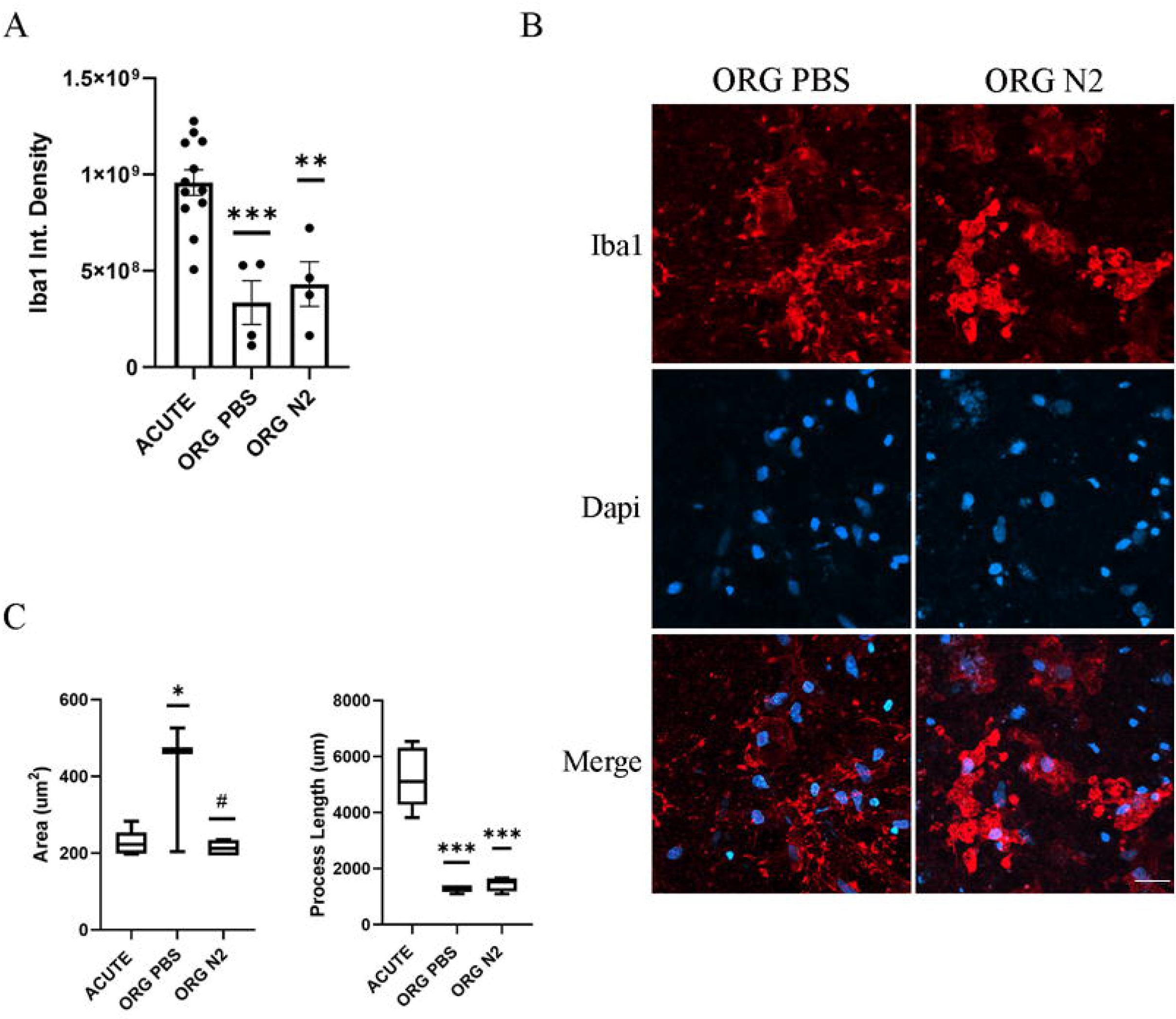
Glioblastoma modifies the human myeloid microenvironment. (**A-B**) Relative quantification of Iba1 density in acute and organotypic human tissue slices injected with PBS or N2 GFP+ cells (p=0.0417, one-way anova, Tukey’s multiple comparisons test, df=12), and representative confocal imaging. Iba1, red. Nuclei were stained by using Dapi (blue). Scale bar 20 µm. (**C**) Morphometric measurements of area and processes length for Iba1+ cells from confocal imaging of acute and organotypic tissues injected with PBS or N2 GFP+ cells. Ordinary One way Anova followed by Tukey’s multiple comparisons test, *,^#^p≤0.05 ***p≤0.0001.

Altogether these results show there are potential modifications involving resident astrocytes and myeloid cells in both the organotypic models. This may support the spreading of GBM. Among the molecular features, Cx43 appears to be fine-tuned in the host tissue of both the experimental models.

### 3.6 Cx43 hemichannels may have a role in GBM progression

To dissect how the multifunctional Cx43 protein may work in the GBM spreading, we treated GBM cells with GAP-19, a selective blocker for Cx43 hemichannels, either in culture or in co-culture with brain tissue.

The scratching test revealed that the GAP19 increases the migratory ability of the GBM cells at 4h, with no effect at 24h compared to the CTR group (unpaired t’ student test, p=0,0014, t=4,054; df=13) (**Fig. 8A-B**). GL261 cells responded with an upregulation of Cx43 protein at 4h, and this effect was lost after 24h (CTR=1.477±0.145; GAP19 4h=2.426±0.714; GAP19 24h= 0.867±0.115; #p= 0.014, one-way anova, Tukey’s multiple comparisons test, df=12) (**Fig 8C-D**). No morphological changes were observed with the treatment (**Fig. 8D**).

**Fig. 8.**
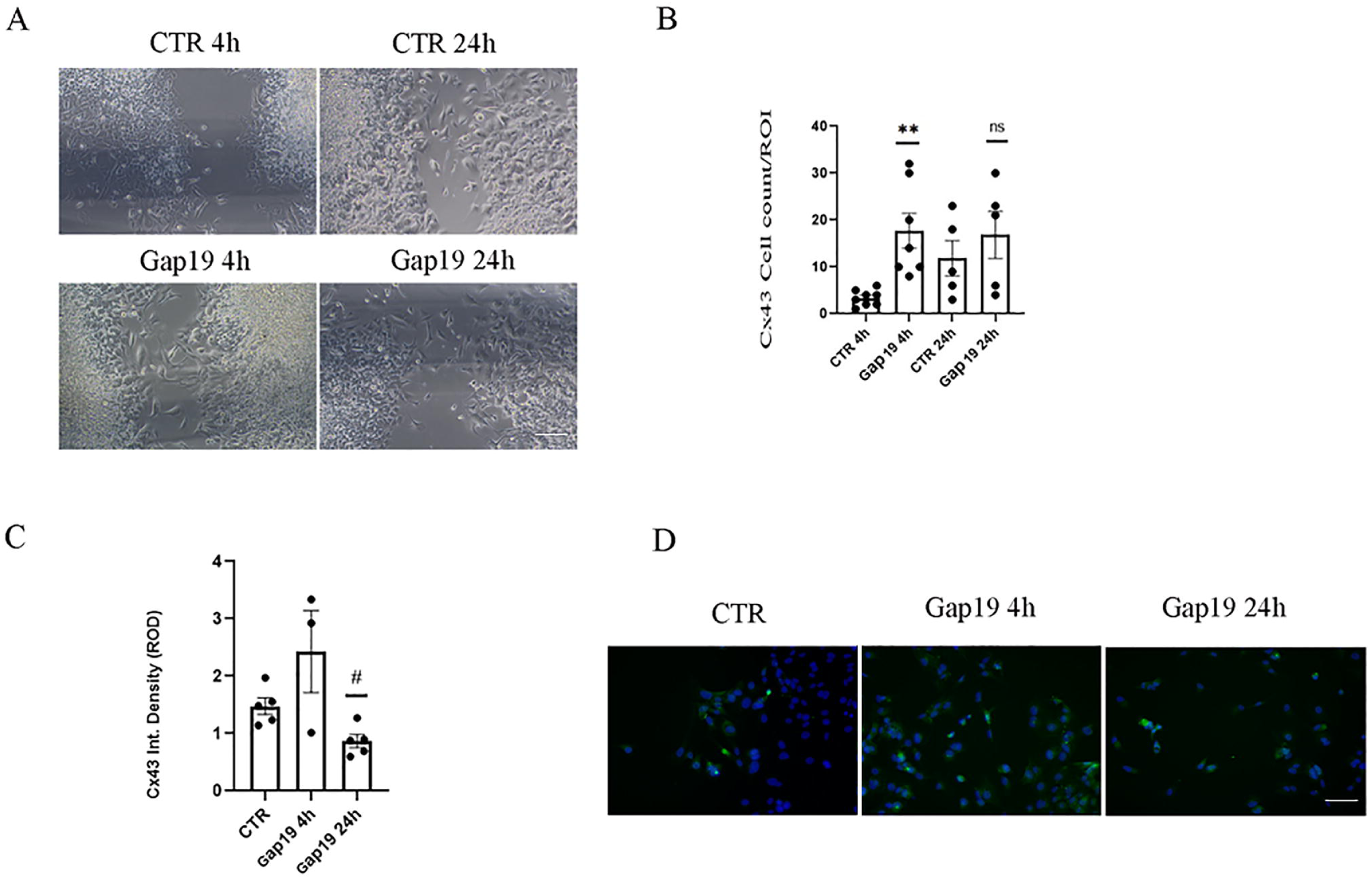
Cx43 protein expression in mouse GL261 cells, and GAP19 effect. **(A-B)** Representative imaging for Cx43 expression and dynamics in GL261 cells culture in CTR conditions or under treatment with GAP19 200 µM at 4 and 24h, and relative quantitation. Cx43, green; DAPI staining for nuclei, blue. Scale bar 50 µm. **(C-D)** Scratching test quantitation for migration ability of GL261 cells cultures in CTR conditions or under treatment with GAP19 200 µM at 4 and 24h, and representative images. Scale bar 50 µm.

On the contrary, using time-lapse microscopy, we found that the eGFP-GBM cells were longitudinally aligned and polarized from DIV 4 to DIV 7 under chronic treatment in culture and co-culture with human tissue (**Fig 9 A-B**). The circularity index of the GFP+ cell bodies in co-culture with brain tissue was significantly lower under GAP19 treatment (p=0.0006, p=0.1921, ordinary one way Anova, Tukey’s multiple comparisons’ test, F (2, 12) = 14.81) **(Fig 9 B-C)**. Both eGFP-GBM and resident cells appeared to display an increase in Cx43 expression at the membrane (**Fig. 9D**).

**Fig. 9.**
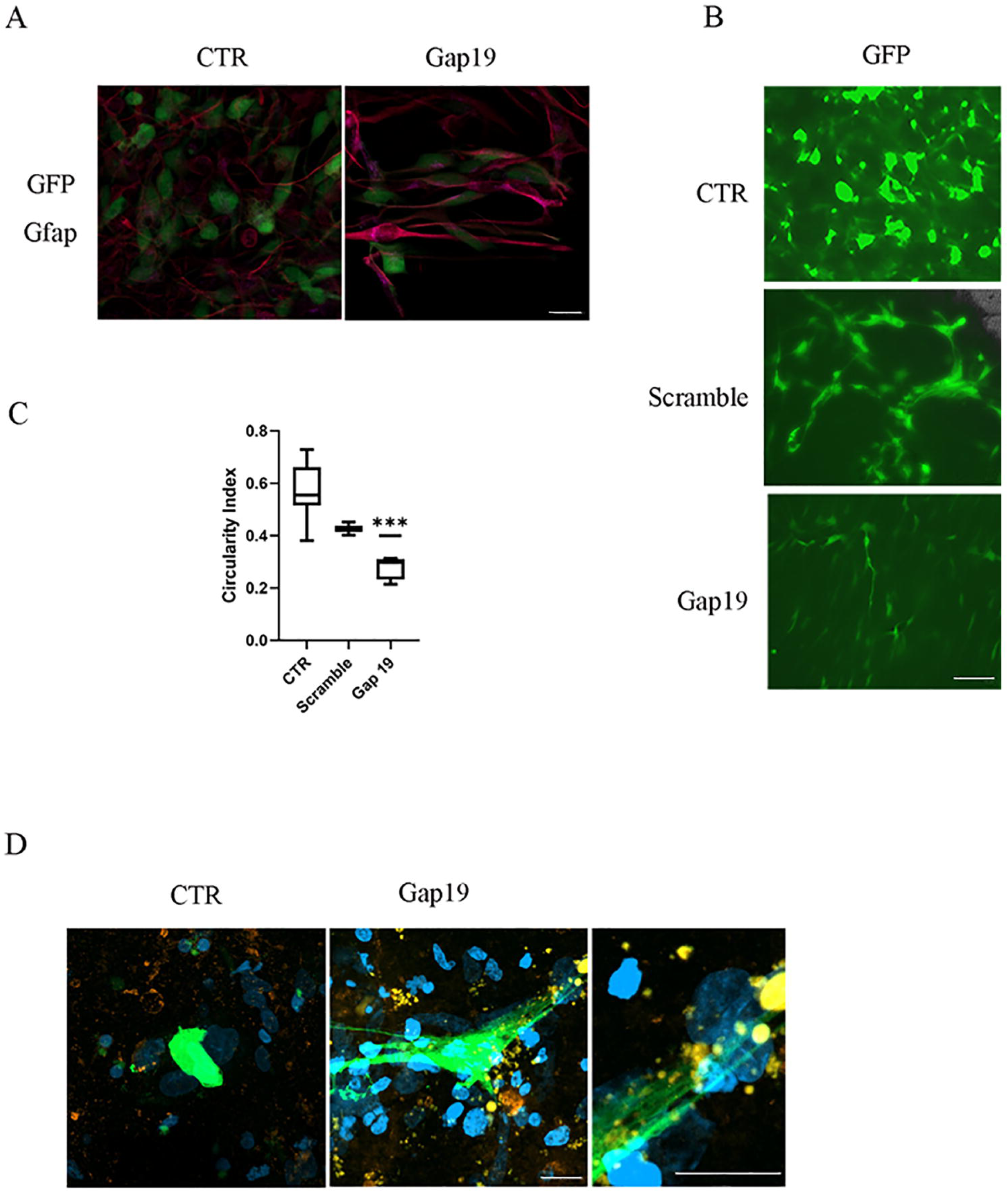
Cx43 protein expression in mouse and human models, and GAP19 effect. (**A**) Representative imaging for GFP+ N2 cells in CTR conditions or under treatment with GAP19 200 µM. (GFP, green; GFAP, pink). Scale bar 20 µm. **(B)** Representative imaging for GFP+ N2 cells in co-culture with human peritumoral organotypic slices, in CTR conditions or under treatment with scramble peptide or GAP19 200 µM. (C) N2 GFP+cells morphometry in co-culture with human peritumoral organotypic slices, in CTR conditions or under treatment with GAP19 200 µM (Ordinary one way Anova, Tukey’s multiple comparisons’ test, p=0.0006, F (2, 12) = 14.81). (**D**) Representative immunofluorescence for Cx43 (orange) of GFP+ N2 cells in co-culture with human peritumoral organotypic slices, in CTR conditions or under treatment with GAP19 200 µM, and detail. Nuclei were stained by Dapi. Scale bar: 20 µm and 16 um, respectively.

Altogether, these data suggest that the blockade of Cx43 hemichannels has a timely response, increases the migratory ability of the GBM cells, and may induce a quick compensatory mobilization of Cx43 reservoirs. Under chronic treatment, Cx43 expression increases in the human GBM microenvironment, indicating a conserved response across the species [21]. Human primary GBM cells appeared to be very sensitive to the GAP19 treatment by changing their shape and polarization.

## 4 Discussion

GBM is a major challenge for research among brain diseases. The high intra- and intertumoral heterogeneity is one of the main barriers, impairing the translational value of rodent experimental models. However, organotypic slice culture enables the study of the human tissue at 3D cellular and small network level, overcoming the temporal limits of the fresh tissue [52]. In this work, we characterized the glial components of the human tumor and peritumoral tissue, both in acute ex vivo and organotypic conditions, followed by the injection of primary GBM cells. We set up a parallel model for the mouse GBM brain to verify its potential translatability. Tissues obtained from patients are always limited in extension. Having a parallel GBM mouse model offering the whole brain could thus be useful to predict the anatomical behavior of the tumor. Despite structural and functional modifications of the neuroglial networks, the human and mouse organotypic slices culture retained the vitality for one week, shedding light on GBM biology, its microenvironment, and offering hints to fight this incurable disease.

The characterization of the fresh human tumor and peritumoral HGG revealed that the tumor core was entangled with myeloid cells. An abundance of macrophages at the tumor core is a hallmark of poor prognosis. Infiltrating macrophages are recruited at the tumor core in the advanced phase of the disease, indicative of cancer recurrence at a short distance [5,53]. Predicting the anatomical and biological behavior of GBM in the brain peritumoral area would be pivotal for planning clinical follow-up, selecting second-line treatments, and improving the quality of life for patients. Myeloid cells in the peritumoral zone of astrocytoma showed a diverse morphology, with a smaller area and longer processes compared to the tumor core, similarly to microglia cells. Rather, the peritumoral microenvironment was characterized by profound changes. Cx43 protein level was upregulated, as well as the CD44 protein. Astrocytic Cx43 in the peritumoral microenvironment affects the microglial functions and has been implicated in gliomagenesis, tumor progression, and treatment resistance [33,54,55]. The ECM receptor CD44 might coordinate with glial activities in the tumor microenvironment and its expression at the GBM periphery could provide a reliable biomarker for tumor invasiveness [56–58].

We found an increase in Cx43+ cells and Cx43 expression in the host tissue following the infiltration of GBM in human and mouse models, compared to the organotypic control group. Accordingly, an upregulation of Cx43 has been detected at the tumor edge, potentially to promote tumor invasion [14,59]. Cx43-KO does not induce any compensation of other connexins such as Cx30 [60], implying that the Cx43 response to GBM may constitute a conserved mechanism across the two species.

Cx43 expression changes were related to both the tumor cells and the resident tissue. Accordingly, Cx43 should facilitate the detachment of GBM cells from the tumor core and enhance the movements in the parenchyma [33], while reactive astrocytes are known to overexpress Cx43 following brain injury and inflammation [61,62].The increase in Cx43+ cell count in the adult brain slice system may result from the injected tumor cells, which are capable of proliferating as demonstrated by ki67 staining. However, the expression changes of Cx43 could be due to an increase in the transcriptional activity or changes in its subcellular distribution [21,63]. Indeed, Cx43 expression is dynamic, and its localization shifts from being cytoplasmic to the membrane-bound. In breast cancer, the nuclear translocation and localization of Cx43 have been observed [49]. In GBM stem cells, non-junctional Cx43 interacts with the microtubule net for cell maintenance and tumorigenicity [64]. Fresh brain slices and 2D cultures of murine GBM cells reported a Cx43 distribution ratio (cytoplasmic/membrane) close to 1 and 0, respectively, demonstrating that Cx43 protein levels in brain tissue are almost in equilibrium among the cytoplasm and membrane compartments, while GBM cells express Cx43 at low levels and membrane-bound. The coculture of GBM cells and brain slices at 3 dpi showed an increase of the “cytoplasmic” term of the ratio, suggesting a mobilization for protein biogenesis. Cx43 cellular expression is finely and continuously regulated, suggesting the importance for the astrocytic-like cells, including the GBM cells. Accordingly, the blockage of the hemi-channels by GAP19 induced a further upregulation of the Cx43 protein, apparently with a shift toward the membrane localization in the human organotypic model. In addition, human primary GBM cells modified their morphology and polarization under Cx43 hemichannel blockage. The e-GFP labeled GBM cells appeared elongated with fusiform bidirectional processes and polarized toward the same direction, with a palisade aspect to remind the migratory phenotype of their potential ancestors [65]. Accordingly, astrocytes shape and polarity are strongly modified during cancer development, and this is linked to changes in ECM composition, soluble factors, and cytoskeletal forces [66,67]. Blockage of the Cx43 hemichannels could lead to the lack of appropriate, in vivo, cues from the microenvironment, affecting GBM cells morphology and inducing the oncostreams, phenomena of collective migration [68].

GL261 cells showed an increase in migratory ability in the same condition.

In conclusion, the present study provides insights into GBM biology and microenvironment, enabling the use of human and mouse organotypic brain slice cultures as valuable tools for basic neuroscience and GBM disease modeling [69]. Interestingly, the glial Cx43 seems to be a hub in response to GBM progression, which points to Cx43-targeted therapy. Mimicking a better in vitro microenvironment of malignant glioma, including the vasculature or ECM soluble factors, will provide opportunities for patient-specific drug screening, prompting knowledge and further studies [70].

## Acknowledgements and Funding

This work was supported by the Italian Society of Anatomy and Histology, Fellowship 2022 to A.V; Fondazione Bartolo Longo III Millennio to G.C.; the European Innovation Council (EIC)-Pathfinder Programme (THOR; grant number 101099719; to M.P., C.D.L., and A.V.); #NEXTGENERATIONEU (NGEU) and the Ministry of University and Research (MUR), National Recovery and Resilience Plan (NRRP), project MNESYS (PE0000006) to M.P. and G.C., project SmarthyVision (P2022JRBMB) to A.V.; the ANR (AstroXCite) to N.R.

## Author Contributions

A.V. performed the experiments, formal analysis, and writing of the original draft; G.M. contributed to experimental part, data collection, formal analysis, and editing of the original draft. C.D.L and J.M., contributed to the experimental part and reviewing of the original draft. An.V., L.R., A.E. contributed to the experimental part and figures preparation. G.C. contributed to conceptualization and reviewing the original draft. G.H. and J.P. provided the human samples and contributed to data collection, conceptualization and reviewing the original draft. A.V., N.R. and M.P. contributed to funding acquisition, supervision, writing, reviewing and editing of the original draft. All the authors approved the manuscript.

## Data Availability

The data that support the findings of this study are available under reasonable request to the corresponding author.

## Ethics Approval

Animal experiments were conducted with approval from the Animal Care and Use Committee under the authorization no. 268/2012-B from the Italian Minister of Health.

## Patient Consent

This study was approved by the Comité d’Evaluation et d’Ethique de l’INSERM - IRB00003888 (approval n°21-864). Subjects’ consent was obtained according to the Declaration of Helsinki.

## References

[1] Farin A, Suzuki S O, Weiker M, Goldman J E, Bruce J N and Canoll P 2006 Transplanted glioma cells migrate and proliferate on host brain vasculature: A dynamic analysis Glia 53 799–808

[2] De Luca C, Virtuoso A, Papa M, Certo F, Barbagallo G M V and Altieri R 2022 Regional Development of Glioblastoma: The Anatomical Conundrum of Cancer Biology and Its Surgical Implication Cells 11 1349

[3] Seker-Polat F, Pinarbasi Degirmenci N, Solaroglu I and Bagci-Onder T 2022 Tumor Cell Infiltration into the Brain in Glioblastoma: From Mechanisms to Clinical Perspectives Cancers (Basel) 14 443

[4] Gupta R K, Niklasson M, Bergström T, Segerman A, Betsholtz C and Westermark B 2024 Tumor-specific migration routes of xenotransplanted human glioblastoma cells in mouse brain Sci Rep 14 864

[5] Virtuoso A, De Luca C, Cirillo G, Riva M, Romano G, Bentivegna A, Lavitrano M, Papa M and Giovannoni R 2022 Tumor Microenvironment and Immune Escape in the Time Course of Glioblastoma Mol Neurobiol 59 6857–73

[6] Kirschenbaum D, Xie K, Ingelfinger F, Katzenelenbogen Y, Abadie K, Look T, Sheban F, Phan T S, Li B, Zwicky P, Yofe I, David E, Mazuz K, Hou J, Chen Y, Shaim H, Shanley M, Becker S, Qian J, Colonna M, Ginhoux F, Rezvani K, Theis F J, Yosef N, Weiss T, Weiner A and Amit I 2024 Time-resolved single-cell transcriptomics defines immune trajectories in glioblastoma Cell 187 149–165.e23

[7] Mettang M, Meyer-Pannwitt V, Karpel-Massler G, Zhou S, Carragher N O, Föhr K J, Baumann B, Nonnenmacher L, Enzenmüller S, Dahlhaus M, Siegelin M D, Stroh S, Mertens D, Fischer-Posovszky P, Schneider E M, Halatsch M-E, Debatin K-M and Westhoff M-A 2018 Blocking distinct interactions between Glioblastoma cells and their tissue microenvironment: A novel multi-targeted therapeutic approach Sci Rep 8 5527

[8] Sun Y, Wang X, Zhang D Y, Zhang Z, Bhattarai J P, Wang Y, Park K H, Dong W, Hung Y-F, Yang Q, Zhang F, Rajamani K, Mu S, Kennedy B C, Hong Y, Galanaugh J, Sambangi A, Kim S H, Wheeler G, Gonçalves T, Wang Q, Geschwind D H, Kawaguchi R, Viaene A N, Helbig I, Kessler S K, Hoke A, Wang H, Xu F, Binder Z A, Isaac Chen H, Pai E L-L, Stone S, Nasrallah M P, Christian K M, Fuccillo M, Toni N, Wu Z, Cheng H-J, O’Rourke D M, Ma M, Ming G-L and Song H 2025 Brain-wide neuronal circuit connectome of human glioblastoma Nature 641 222–31

[9] Venkataramani V, Tanev D I, Strahle C, Studier-Fischer A, Fankhauser L, Kessler T, Körber C, Kardorff M, Ratliff M, Xie R, Horstmann H, Messer M, Paik S P, Knabbe J, Sahm F, Kurz F T, Acikgöz A A, Herrmannsdörfer F, Agarwal A, Bergles D E, Chalmers A, Miletic H, Turcan S, Mawrin C, Hänggi D, Liu H-K, Wick W, Winkler F and Kuner T 2019 Glutamatergic synaptic input to glioma cells drives brain tumour progression Nature 573 532–8

[10] Virtuoso A, Giovannoni R, De Luca C, Gargano F, Cerasuolo M, Maggio N, Lavitrano M and Papa M 2021 The Glioblastoma Microenvironment: Morphology, Metabolism, and Molecular Signature of Glial Dynamics to Discover Metabolic Rewiring Sequence Int J Mol Sci 22 3301

[11] Resende F F B, Bai X, Del Bel E A, Kirchhoff F, Scheller A and Titze-de-Almeida R 2016 Evaluation of TgH(CX3CR1-EGFP) mice implanted with mCherry-GL261 cells as an in vivo model for morphometrical analysis of glioma-microglia interaction BMC Cancer 16 72

[12] Henrik Heiland D, Ravi V M, Behringer S P, Frenking J H, Wurm J, Joseph K, Garrelfs N W C, Strähle J, Heynckes S, Grauvogel J, Franco P, Mader I, Schneider M, Potthoff A-L, Delev D, Hofmann U G, Fung C, Beck J, Sankowski R, Prinz M and Schnell O 2019 Tumor-associated reactive astrocytes aid the evolution of immunosuppressive environment in glioblastoma Nat Commun 10 2541

[13] Lin C-M, Yu C-F, Huang H-Y, Chen F-H, Hong J-H and Chiang C-S 2019 Distinct Tumor Microenvironment at Tumor Edge as a Result of Astrocyte Activation Is Associated With Therapeutic Resistance for Brain Tumor Front Oncol 9 307

[14] McCutcheon S and Spray D C 2022 Glioblastoma-Astrocyte Connexin 43 Gap Junctions Promote Tumor Invasion Mol Cancer Res 20 319–31

[15] Spalletti C, Scalera M, Mori E, Haddad S, Mainardi M, Cangi D, Pillai V, Parmigiani E, Landi S, Caleo M and Vannini E 2025 Inhibitory circuit dysfunction as a potential contributor to cortical reorganization in Glioblastoma progression Neurobiol Dis 213 106997

[16] Meyer J, Yu K, Luna-Figueroa E, Deneen B and Noebels J 2024 Glioblastoma disrupts cortical network activity at multiple spatial and temporal scales Nat Commun 15 4503

[17] Hausmann D, Hoffmann D C, Venkataramani V, Jung E, Horschitz S, Tetzlaff S K, Jabali A, Hai L, Kessler T, Azoŕin D D, Weil S, Kourtesakis A, Sievers P, Habel A, Breckwoldt M O, Karreman M A, Ratliff M, Messmer J M, Yang Y, Reyhan E, Wendler S, Löb C, Mayer C, Figarella K, Osswald M, Solecki G, Sahm F, Garaschuk O, Kuner T, Koch P, Schlesner M, Wick W and Winkler F 2023 Autonomous rhythmic activity in glioma networks drives brain tumour growth Nature 613 179–86

[18] Díaz-Fernández B, Henao-Herreno D, Nieto J, Evstratova A, Cases-Cunillera S, Deboeuf L, Roux A, Dezamis E, Zanello M, Mathon B, Karachi C, Carpentier A, Varlet P, Pallud J, Capelle L, Alvarado-Rojas C, Le Van Quyen M and Huberfeld G 2025 Differentiation of tumor versus peritumoral cortex in gliomas by intraoperative electrocorticography Neuro Oncol 27 1758–71

[19] Brandao M, Simon T, Critchley G and Giamas G 2019 Astrocytes, the rising stars of the glioblastoma microenvironment Glia 67 779–90

[20] Guillebaud F, Barbot M, Barbouche R, Brézun J-M, Poirot K, Vasile F, Lebrun B, Rouach N, Dallaporta M, Gaige S and Troadec J-D 2020 Blockade of Glial Connexin 43 Hemichannels Reduces Food Intake Cells 9 2387

[21] Kotini M, Barriga E H, Leslie J, Gentzel M, Rauschenberger V, Schambony A and Mayor R 2018 Gap junction protein Connexin-43 is a direct transcriptional regulator of N-cadherin in vivo Nat Commun 9 3846

[22] Delvaeye T, Vandenabeele P, Bultynck G, Leybaert L and Krysko D V 2018 Therapeutic Targeting of Connexin Channels: New Views and Challenges Trends Mol Med 24 1036–53

[23] Tamborini M, Ribecco V, Stanzani E, Sironi A, Tambalo M, Franzone D, Florio E, Fraviga E, Saulle C, Gagliani M C, Pizzocri M, Mattioli M, Cortese K, Jiang J X, Martano G, Politi L S, Riva M, Pessina F, Pozzi D, Lodato S, Passoni L and Matteoli M 2025 Extracellular Vesicles Released by Glioblastoma Cancer Cells Drive Tumor Invasiveness via Connexin-43 Gap Junctions Neuro Oncol noaf013

[24] Orellana J A, Froger N, Ezan P, Jiang J X, Bennett M V L, Naus C C, Giaume C and Sáez J C 2011 ATP and glutamate released via astroglial connexin 43 hemichannels mediate neuronal death through activation of pannexin 1 hemi-channels J Neurochem 118 826–40

[25] Pannasch U, Vargová L, Reingruber J, Ezan P, Holcman D, Giaume C, Syková E and Rouach N 2011 Astroglial networks scale synaptic activity and plasticity Proc Natl Acad Sci U S A 108 8467–72

[26] Medina-Ceja L, Salazar-Sánchez J C, Ortega-Ibarra J and Morales-Villagrán A 2019 Connexins-Based Hemichannels/Channels and Their Relationship with Inflammation, Seizures and Epilepsy Int J Mol Sci 20 5976

[27] Chever O, Lee C-Y and Rouach N 2014 Astroglial connexin43 hemichannels tune basal excitatory synaptic transmission J Neurosci 34 11228–32

[28] Schumacher S, Tahiri H, Ezan P, Rouach N, Witschas K and Leybaert L 2024 Inhibiting astrocyte connexin-43 hemichannels blocks radiation-induced vesicular VEGF-A release and blood-brain barrier dysfunction Glia 72 34–50

[29] Guo A, Zhang H, Li H, Chiu A, García-Rodríguez C, Lagos C F, Sáez J C and Lau C G 2022 Inhibition of connexin hemichannels alleviates neuroinflammation and hyperexcitability in temporal lobe epilepsy Proc Natl Acad Sci U S A 119 e2213162119

[30] Donati V, Di Pietro C, Persano L, Rampazzo E, Panarelli M, Cambria C, Selimi A, Manfreda L, de Oliveira do Rêgo A G, La Sala G, Sprega C, Calistri A, Ciubotaru C D, Yang G, Zonta F, Antonucci F, Marazziti D and Mammano F 2025 Connexin hemichannel blockade by abEC1.1 disrupts glioblastoma progression, suppresses invasiveness, and reduces hyperexcitability in preclinical models Cell Commun Signal 23 391

[31] Murphy S F, Varghese R T, Lamouille S, Guo S, Pridham K J, Kanabur P, Osimani A M, Sharma S, Jourdan J, Rodgers C M, Simonds G R, Gourdie R G and Sheng Z 2016 Connexin 43 Inhibition Sensitizes Chemoresistant Glioblastoma Cells to Temozolomide Cancer Res 76 139–49

[32] Uzu M, Sin W C, Shimizu A and Sato H 2018 Conflicting Roles of Connexin43 in Tumor Invasion and Growth in the Central Nervous System Int J Mol Sci 19 1159

[33] Sin W C, Aftab Q, Bechberger J F, Leung J H, Chen H and Naus C C 2016 Astrocytes promote glioma invasion via the gap junction protein connexin43 Oncogene 35 1504–16

[34] Szatmári T, Lumniczky K, Désaknai S, Trajcevski S, Hídvégi E J, Hamada H and Sáfrány G 2006 Detailed characterization of the mouse glioma 261 tumor model for experimental glioblastoma therapy Cancer Sci 97 546–53

[35] Ciraku L, Moeller R A, Esquea E M, Gocal W A, Hartsough E J, Simone N L, Jackson J G and Reginato M J 2021 An Ex Vivo Brain Slice Model to Study and Target Breast Cancer Brain Metastatic Tumor Growth J Vis Exp

[36] Virtuoso A, Galanis C, Lenz M, Papa M and Vlachos A 2024 Regional Microglial Response in Entorhino-Hippocampal Slice Cultures to Schaffer Collateral Lesion and Metalloproteinases Modulation Int J Mol Sci 25 2346

[37] Dossi E, Blauwblomme T, Nabbout R, Huberfeld G and Rouach N 2014 Multielectrode array recordings of human epileptic postoperative cortical tissue J Vis Exp e51870

[38] Maugeri G, D’Amico A G, Amenta A, Saccone S, Federico C, Reibaldi M, Russo A, Bonfiglio V, Avitabile T, Longo A and D’Agata V 2020 Protective effect of PACAP against ultraviolet B radiation-induced human corneal endothelial cell injury Neuropeptides 79 101978

[39] Miceli M, Dell’Aversana C, Russo R, Rega C, Cupelli L, Ruvo M, Altucci L and Chambery A 2016 Secretome profiling of cytokines and growth factors reveals that neuro-glial differentiation is associated with the down-regulation of Chemokine Ligand 2 (MCP-1/CCL2) in amniotic fluid derived-mesenchymal progenitor cells Proteomics 16 674–88

[40] Marques-Torrejon M A, Gangoso E and Pollard S M 2018 Modelling glioblastoma tumour-host cell interactions using adult brain organotypic slice coculture Dis Model Mech 11 dmm031435

[41] Gump J M and Dowdy S F 2007 TAT transduction: the molecular mechanism and therapeutic prospects Trends Mol Med 13 443–8

[42] Lissoni A, Wang N, Nezlobinskii T, De Smet M, Panfilov A V, Vandersickel N, Leybaert L and Witschas K 2020 Gap19, a Cx43 Hemichannel Inhibitor, Acts as a Gating Modifier That Decreases Main State Opening While Increasing Substate Gating Int J Mol Sci 21 7340

[43] De Bock M, De Smet M, Verwaerde S, Tahiri H, Schumacher S, Van Haver V, Witschas K, Steinhäuser C, Rouach N, Vandenbroucke R E and Leybaert L 2022 Targeting gliovascular connexins prevents inflammatory blood-brain barrier leakage and astrogliosis JCI Insight 7 e135263

[44] Dossi E, Zonca L, Pivonkova H, Milior G, Moulard J, Vargova L, Chever O, Holcman D and Rouach N 2024 Astroglial gap junctions strengthen hippocampal network activity by sustaining afterhyperpolarization via KCNQ channels Cell Rep 43 114158

[45] Cirillo G, Colangelo A M, Bianco M R, Cavaliere C, Zaccaro L, Sarmientos P, Alberghina L and Papa M 2012 BB14, a Nerve Growth Factor (NGF)-like peptide shown to be effective in reducing reactive astrogliosis and restoring synaptic homeostasis in a rat model of peripheral nerve injury Biotechnol Adv 30 223–32

[46] Croft C L and Noble W 2018 Preparation of organotypic brain slice cultures for the study of Alzheimer’s disease F1000Res 7 592

[47] Bissen D, Kracht M K, Foss F and Acker-Palmer A 2022 Expansion microscopy of mouse brain organotypic slice cultures to study protein distribution STAR Protoc 3 101507

[48] Ylagan L R and Quinn B 1997 CD44 expression in astrocytic tumors Mod Pathol 10 1239–46

[49] Pilarczyk G, Papenfuß F, Bestvater F and Hausmann M 2019 Spatial Arrangements of Connexin43 in Cancer Related Cells and Re-Arrangements under Treatment Conditions: Investigations on the Nano-Scale by Super-Resolution Localization Light Microscopy Cancers (Basel) 11 301

[50] Humpel C 2015 Organotypic brain slice cultures: A review Neuroscience 305 86–98

[51] Harwood D S L, Pedersen V, Bager N S, Schmidt A Y, Stannius T O, Areškevičiūtė A, Josefsen K, Nørøxe D S, Scheie D, Rostalski H, Lü M J S, Locallo A, Lassen U, Bagger F O, Weischenfeldt J, Heiland D H, Vitting-Seerup K, Michaelsen S R and Kristensen B W 2024 Glioblastoma cells increase expression of notch signaling and synaptic genes within infiltrated brain tissue Nat Commun 15 7857

[52] Bak A, Koch H, van Loo K M J, Schmied K, Gittel B, Weber Y, Ort J, Schwarz N, Tauber S C, Wuttke T V and Delev D 2024 Human organotypic brain slice cultures: a detailed and improved protocol for preparation and long-term maintenance J Neurosci Methods 404 110055

[53] Shojaee P, Weinholtz E, Schaadt N S, Feuerhake F and Hatzikirou H 2025 Biopsy location and tumor-associated macrophages in predicting malignant glioma recurrence using an in-silico model NPJ Syst Biol Appl 11 3

[54] Sin W-C, Crespin S and Mesnil M 2012 Opposing roles of connexin43 in glioma progression Biochim Biophys Acta 1818 2058–67

[55] Wang L, Peng Y, Peng J, Shao M, Ma L, Zhu Z, Zhong G, Xia Z and Huang H 2018 Tramadol attenuates the sensitivity of glioblastoma to temozolomide through the suppression of Cx43-mediated gap junction intercellular communication Int J Oncol 52 295–304

[56] Inoue A, Ohnishi T, Nishikawa M, Watanabe H, Kusakabe K, Taniwaki M, Yano H, Ohtsuka Y, Matsumoto S, Suehiro S, Yamashita D, Shigekawa S, Takahashi H, Kitazawa R, Tanaka J and Kunieda T 2023 Identification of CD44 as a Reliable Biomarker for Glioblastoma Invasion: Based on Magnetic Resonance Imaging and Spectroscopic Analysis of 5-Aminolevulinic Acid Fluorescence Biomedicines 11 2369

[57] Du Z, Wang Y, Liang J, Gao S, Cai X, Yu Y, Qi Z, Li J, Xie Y and Wang Z 2022 Association of glioma CD44 expression with glial dynamics in the tumour microenvironment and patient prognosis Comput Struct Biotechnol J 20 5203–17

[58] Lubanska D, Alrashed S, Mason G T, Nadeem F, Awada A, DiPasquale M, Sorge A, Malik A, Kojic M, Soliman M A R, deCarvalho A C, Shamisa A, Kulkarni S, Marquardt D, Porter L A and Rondeau-Gagné S 2022 Impairing proliferation of glioblastoma multiforme with CD44+ selective conjugated polymer nanoparticles Sci Rep 12 12078

[59] Baklaushev V P, Yusubalieva G M, Tsitrin E B, Gurina O I, Grinenko N P, Victorov I V and Chekhonin V P 2011 Visualization of Connexin 43-positive cells of glioma and the periglioma zone by means of intravenously injected monoclonal antibodies Drug Deliv 18 331–7

[60] Pridham K J, Shah F, Hutchings K R, Sheng K L, Guo S, Liu M, Kanabur P, Lamouille S, Lewis G, Morales M, Jourdan J, Grek C L, Ghatnekar G G, Varghese R, Kelly D F, Gourdie R G and Sheng Z 2022 Connexin 43 confers chemoresistance through activating PI3K Oncogenesis 11 2

[61] Theodoric N, Bechberger J F, Naus C C and Sin W-C 2012 Role of gap junction protein connexin43 in astrogliosis induced by brain injury PLoS One 7 e47311

[62] Colangelo A M, Cirillo G, Lavitrano M L, Alberghina L and Papa M 2012 Targeting reactive astrogliosis by novel biotechnological strategies Biotechnol Adv 30 261–71

[63] Ochalski P A, Sawchuk M A, Hertzberg E L and Nagy J I 1995 Astrocytic gap junction removal, connexin43 redistribution, and epitope masking at excitatory amino acid lesion sites in rat brain Glia 14 279–94

[64] Smyth J W, Guo S, Chaunsali L, O’Rourke L, Dahlka J, Deaver S, Lunski M, Nurmemmedov E, Sontheimer H, Sheng Z, Gourdie R G and Lamouille S 2025 Cytoplasmic connexin43-microtubule interactions promote glioblastoma stem-like cell maintenance and tumorigenicity Cell Death Dis 16 388

[65] John Lin C-C, Yu K, Hatcher A, Huang T-W, Lee H K, Carlson J, Weston M C, Chen F, Zhang Y, Zhu W, Mohila C A, Ahmed N, Patel A J, Arenkiel B R, Noebels J L, Creighton C J and Deneen B 2017 Identification of diverse astrocyte populations and their malignant analogs Nat Neurosci 20 396–405

[66] Purvis E M, O’Donnell J C and Cullen D K 2022 Unique Astrocyte Cytoskeletal and Nuclear Morphology in a Three-Dimensional Tissue-Engineered Rostral Migratory Stream Neuroglia 3 41–60

[67] Etienne-Manneville S 2008 Polarity proteins in glial cell functions Curr Opin Neurobiol 18 488–94

[68] Kang S, Ughetta M E, Zhang J Y, Marallano V J, Sattiraju A, Hannah T, Wahane S, Ramakrishnan A, Estill M, Tsankova N M, Shen L, Tsankov A M, Friedel R H and Zou H 2025 Glioblastoma shift from bulk to infiltrative growth is guided by plexin-B2-mediated microglia alignment in invasive niches Nat Cancer 6 1505–23

[69] Mauro A and Mohamed S 2020 Three dimensional heat and mass transfer in human eye based on porous medium approach International Journal of Heat and Mass Transfer 158 119994

[70] De Luca C and Papa M 2016 Looking Inside the Matrix: Perineuronal Nets in Plasticity, Maladaptive Plasticity and Neurological Disorders Neurochem Res 41 1507–15

